# Detection of Parasites in Microbiomes using Metagenomics

**DOI:** 10.1101/2022.03.27.485979

**Authors:** Philipp Kirstahler, Frank M. Aarestrup, Sünje Johanna Pamp

## Abstract

Despite a yearly death toll of up to one million people due to parasite-related infections, parasites are still neglected in genomics research. While there is progress in the detection of bacteria and viruses using metagenomics in the context of infectious diseases, there are still challenges in metagenomics-based detection of parasites. Here, we implement a workflow for the detection of parasites from metagenomics data. We employ stringent cut off criteria to limit false positive detections. We analysed a total of 7.120 metagenomics samples of which 359 originated from gut microbiomes of livestock (pigs and chicken) from nine countries, and 6.761 from gut microbiomes of humans (adults and infants) from 25 countries. Five parasite-related genera were detected in livestock, of which *Blastocystis* sp. was detected in 71% of all pig herds and *Eimeria* in 83% of all chicken flocks. Distinct gut bacterial taxa were associated with *Blastocystis* sp. abundance in pigs. Nine parasite-related genera were detected in humans. *Blastocystis* sp. subtypes ST1, ST2, and ST3 were detected in all countries, and ST3 was most predominant. A higher overall prevalence of *Blastocystis* sp. was observed in low-income countries as compared to high-income countries, and a higher diversity of *Blastocystis* subtypes (ST1, ST2, ST3, ST4, ST6, ST7, ST8) was detected in high-income countries as compared to low-income countries. The prevalence of *Blastocystis* sp. in infant gut microbiome samples was lower as compared to adults. Overall, metagenomics-based analysis may be a promising tool for parasite detection from complex microbiome samples in clinical and veterinary medicine.

Metagenomics could become the preferred method for parasite detection for a wide range of biological samples. Current parasite detection methods often rely on microscopic examination of the sample or using specific PCR. Metagenomics-based analyses may allow for a faster and more convenient way of detecting parasites in humans and animals, as this approach could serve as a one-for-all untargeted approach for pathogen detection, including bacteria, viruses, and parasites.

## Introduction

Eukaryotic parasites belong to a diverse group of organisms, and there are currently various diagnostic approaches to detect parasites in biological specimens. The most common methods in human and veterinary medicine involve microscopic examination of the sample for parasite eggs or the parasite itself. In addition to these, specific serology and molecular-based assays, for example enzyme-linked immunosorbent assay (ELISA) or polymerase chain reaction (PCR), have been developed (1–5).

Metagenomics is increasingly being used as a tool for pathogen identification in the context of various diseases and for different sample types (6–9). However, only a small fraction of these studies deal with actual parasite infections (10, 11). Most of these studies focus on the detection of a particular single parasite, or the analysis of the sample for their bacterial content using metagenomics while parasites are detected using traditional methods, for example the Kato-Katz procedure and qPCR analysis (12, 13). There are several reasons for why parasites are seldomly analysed using metagenomics so far. First, the majority of human infections with parasites happen in underdeveloped countries, which may make parasite investigations less of a research priority in high-development countries. Second, parasites are not a homogenous group, they are grouped by their way of living and not based on evolutionary similarities. This creates a set of organisms with different genomic properties, from small genomes with around 6 million bases to large genomes with 1.5 billion bases. Third, with less than a thousand available genomes, parasites are one of the least studied groups in genomics. In comparison, there are over 18,000 *Escherichia coli* genomes available at NCBI/EBI/DDBJ. In addition, these parasite genomes are often highly fragmented and contain contaminations with foreign DNA, likely introduced during sample collection and DNA extraction. First attempts to clean parasite genomes for use in metagenomic parasite detection in human samples were performed recently (6, 14).

Here, we developed a workflow for the detection of parasites (for which reference genome sequences exist) from metagenomics shotgun data sets. The workflow involves a removal of potential contaminations from the reference parasite genomes. We applied our approach to a total of 7.120 fecal samples from livestock and humans. We find distinct parasites in the samples and discuss how parasite detection can be improved in the future.

## Results

A total of 7.120 metagenomics samples were assessed in this study. We examined metagenomic data from 359 gut microbiome samples from livestock of which 181 originated from pigs and 178 from chicken from a total of nine European countries (Figure 1A) (15). For each country, between 19 and 21 farms were investigated for pig and chicken respectively. In addition, we examined metagenomic sequencing data from 6.152 gut microbiome samples from human adults and 609 fecal samples from human infants originating from 22 and 4 different countries, respectively. The number of reads for the livestock samples were on average 47 million for chicken (range: 26 to 183 million read-pairs) and 44.4 million for pig samples (range: 3 to 97 million read-pairs). The number of reads for the human gut samples were on average 23.2 million reads (range: 10 to 44 million read-pairs) for adults, and 19.0 million reads (range 10.0 to 38.4 million read-pairs) for infants (Figure 1B). The metagenomic sequencing reads were analysed using kraken 2 and a microbial genome reference database generated using carefully chosen cut-offs to limit false-positive parasite identifications and remove parasite identifications below a defined detection threshold (see Materials & Methods, and Figure 1C).

**Figure 1.**
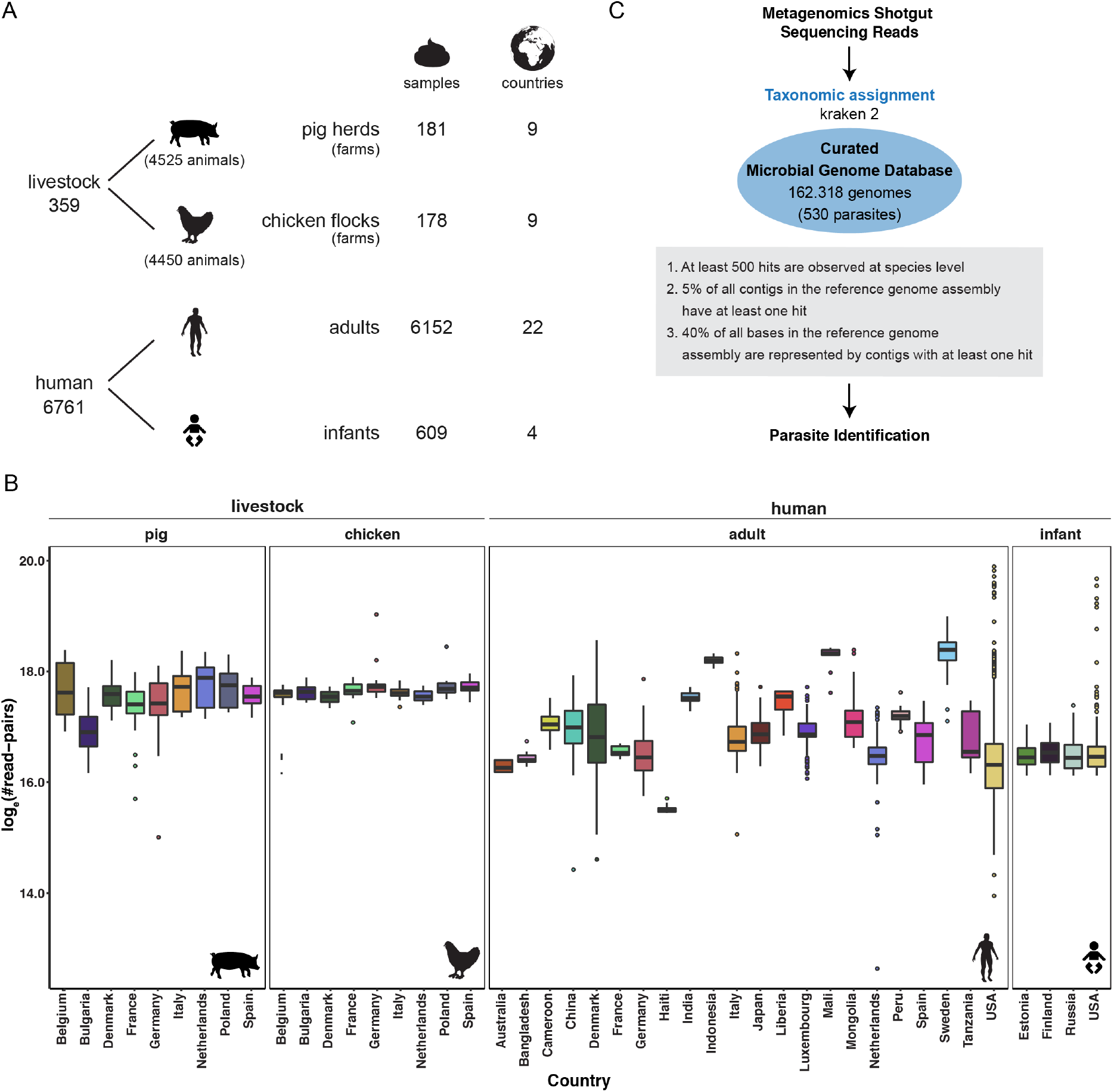
Overview of Data and Workflow for Data Analysis. A) A total of 7120 metagenomics samples were analysed that originated from livestock and human fecal samples. Each livestock sample represents a single pooled sample from a distinct farm from nine European countries. The pooled samples are composed of 25 individual fecal samples from distinct animals at the farm, respectively. B) Logarithmic read counts of each sample arranged by sample type and country. C) Outline of the metagenomics data analysis workflow employed in this study, involving kraken 2 and a reference genome database and distinct quality cut-offs.

### Parasites detected in pig and chicken gut microbiomes

In the pig and chicken gut microbiomes, we detected a total of 47 different parasite-related genera through metagenomic read classification (Figure S1). Upon applying the quality cut-offs (see Methods), five parasite-related genera remained (*Ascaris, Blastocystis, Cryptosporidium, Eimeria, Pristionchus*) of which four parasite-related genera were detected in multiple samples (*Ascaris, Blastocystis, Cryptosporidium, Eimeria*) (Figure 2A).

**Figure 2.**
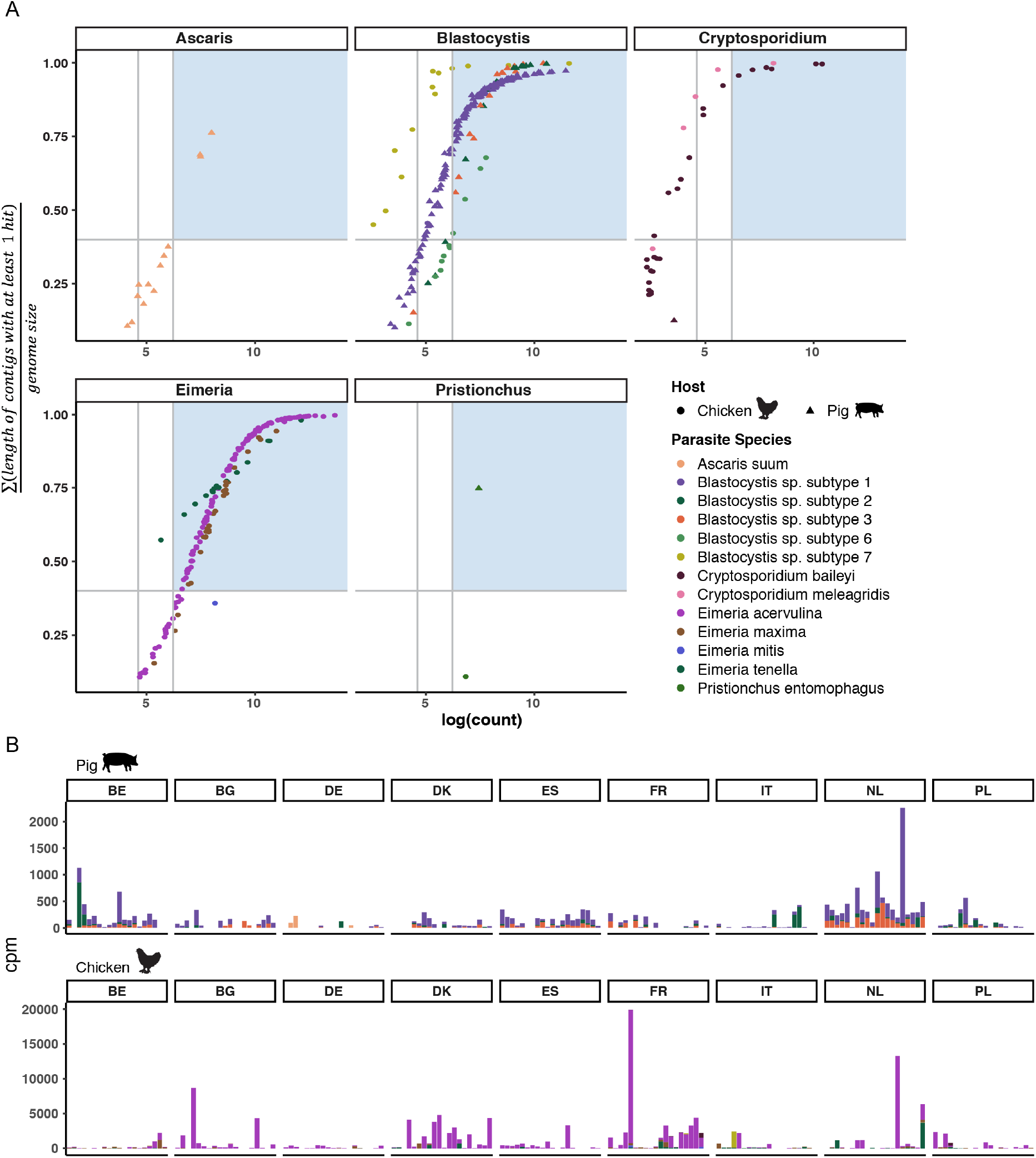
Parasites detected in livestock samples. A) The dot plot depicts parasite-related genera in livestock samples for which at least one parasite-related species/subtype satisfied the cut-off criteria. The data points are coloured according to species and indicated by the host they originate from. The first vertical grey line indicates the 100-read count threshold, and the second line represents the selected cut-off of 500 reads observed at species level. The horizontal grey line indicates the cut-off for the sum of the length of all contigs with hits (0.4). The light blue area indicates the samples that passed the cut-off criteria. B) The bar plot indicates the abundance of parasites detected at individual farms in the different countries. The counts indicate count per million (cpm) to adjust for library size. Blastocystis is predominantly found in pig herds, while chicken flocks are predominantly infected with Eimeria.

*Blastocystis* was detected in 129 (70%) of the pig herds. Pig feces samples from the Netherlands had a higher abundance of *Blastocystis* compared to samples from other countries (Figure 2B). *Blastocystis ST1* was the dominant subtype in pigs in all countries followed by ST3 and ST2 (Figure 2B, and Table S1). Other parasites detected in pig herds were *Ascaris suum* in three pig herds from Germany and *Pristionchus entomophagus* in one pig herd in Poland (Table S2).

*Eimeria* was detected in 148 chicken flocks, accounting for 82% of all chicken samples. Overall, *Eimeria acervulina* was the dominant *Eimeria* species in all countries with the exception of Belgium (Figure 2B). In Belgium, 11 out of 20 flocks were colonized with *Eimeria maxima* (Figure 2B, and Table S3). In other countries, the number of flocks colonized with *Eimeria maxima* ranged between 0 and 20% of the samples. Furthermore, *Eimeria tenella* was present in up to 20% of the flocks in five countries (Table S3). In addition, we identified *Cryptosporidium* (*C. baileyi* and *C. meleagridis*) in seven chicken flocks of which four were located in France. Moreover, we identified *Blastocystis* (ST6 and ST7) in seven flocks in four countries (Table S4).

### Blastocystis and Eimeria are differentially abundant between countries

Pairwise comparison of the abundance of *Blastocystis* between countries revealed elevated levels of *Blastocystis* (genus) and *Blastocystis* sp. subtype ST1, ST2, and ST3 in some countries. For example, *Blastocystis* was significantly higher abundant in pig farms from the Netherlands and Belgium as compared to most other countries (Figure 3). In contrast, *Blastocystis* genus and subtype levels (ST1, ST2, ST3) were significantly lower in Germany, Poland, Bulgaria, and Italy. The lowest abundance of *Blastocystis* was observed in pig herds from Germany for all three subtypes, and for pig farms from Italy for subtype ST2 and ST3. Although some differences were observed, no significant differences for *Eimeria* species in chicken flocks were detected between countries (Figure 3).

**Figure 3.**
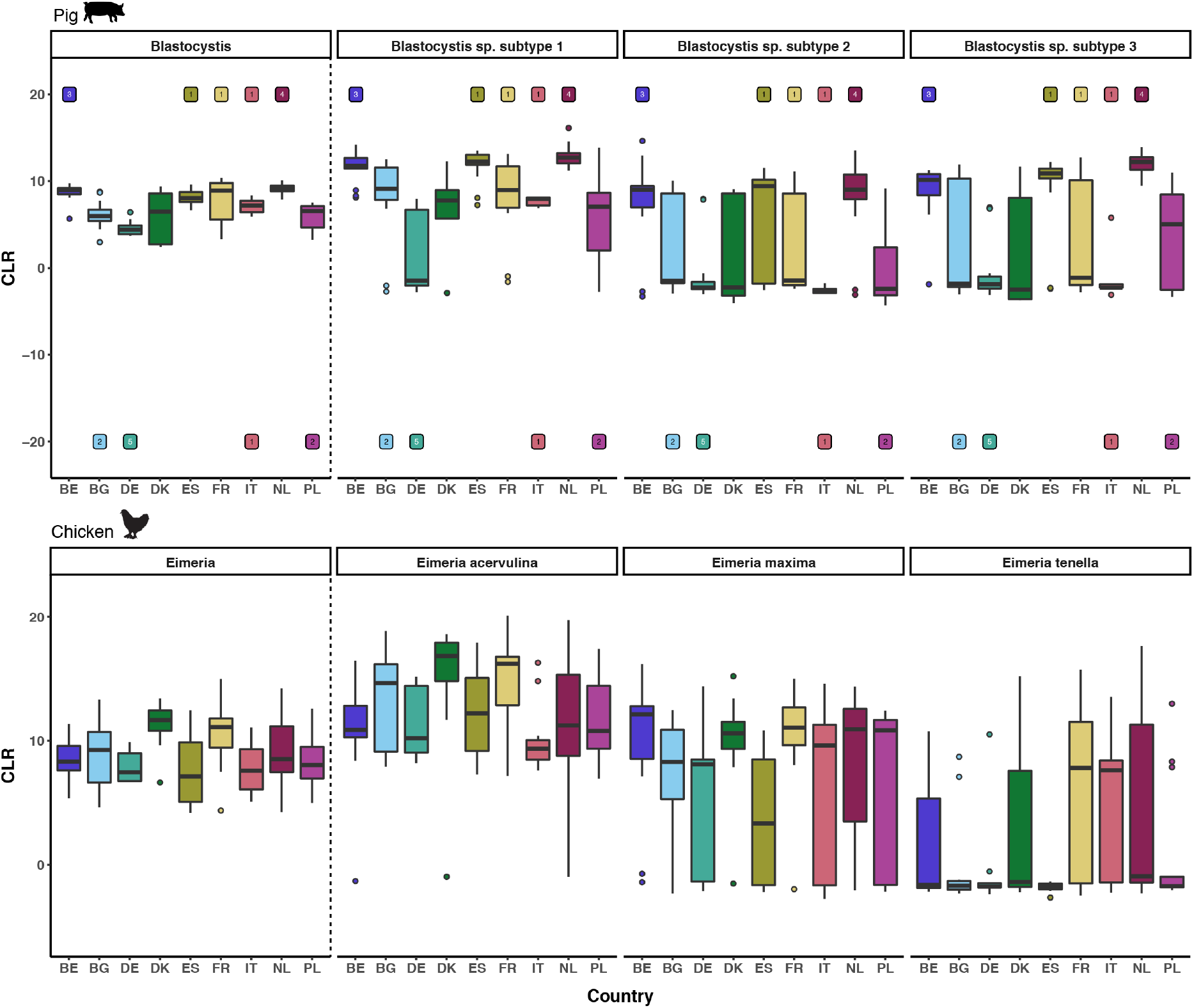
Differential abundance of *Blastocystis* and *Eimeria* between European countries. The boxplot displays centered log ratio (CLR) values for the *Blastocystis* genus and subtypes (ST1, ST2, ST3) and *Eimeria* genus and species. The upper and lower whisker extends from the hinge to the largest and smallest value no further than 1.5 * the IQR from the hinge, respectively. Individually plotted points are outliers. Numeric labels at the top and bottom indicate the instances of significant increased or decreased abundance in pairwise comparisons to other countries.

### Bacteria of the pig gut microbiome positively associated with Blastocystis

To test whether other gut microbiome members in pigs and chicken were associated with the presence of *Blastocystis* and *Eimeria* respectively, we used the definition to classify a sample as positive/negative using the cut-off criteria as described in Materials & Methods. Beside taxa that are part of the *Blastocystis* lineage, we identified a set of bacteria in the pig gut microbiome that were positively associated with the presence of *Blastocystis*. For example, several species and/or strains affiliated with the bacterial genus *Prevotella* (Bacteroidetes) were positively associated with Blastocystis abundance (Figure S2). In addition, taxa of the Firmicutes phylum, including strains affiliated with *Roseburia* and *Ruminococcus* were positively associated with Blastocystis abundance. In chicken we did not identify specific gut bacteria positively associated with *Eimeria*. When we compared *Eimeria*-positive and *Eimeria*-negative gut samples, only taxa related to the *Eimeria* lineage were identified (Figure S3). In neither the pig nor the chicken samples we identified taxa negatively associated with the respective parasites.

### Parasites detected in human adult gut microbiomes

In the gut microbiome samples from human adults, we identified 9 parasite-related genera (Figure 4 and Figure S4). The most prevalent parasite was *Blastocystis* which was identified in 870 samples from all 22 countries investigated here. In addition, *Giardia intestinalis* was detected in 18 samples, of which 8 were collected in Germany. Two types of nematodes (roundworms) were identified: *Ascaris suum* in 11 samples and *Necator americanus* in 12 samples. Most of the detected *Ascaris suum* and *Necator americanus* cases were from a study investigating soil-transmitted parasitic-worms in Indonesia and Liberia (13). Other detected parasites were: *Clonorchis sinensis* (Chinese liver fluke) in two samples from China, *Schistosoma mansoni* (a blood fluke) in one sample each from Cameroon and Mali, and *Strongyloides stercoralis* (threadworm) in one sample from Cameroon (Figure 4). A detailed overview is provided in Table S5.

**Figure 4.**
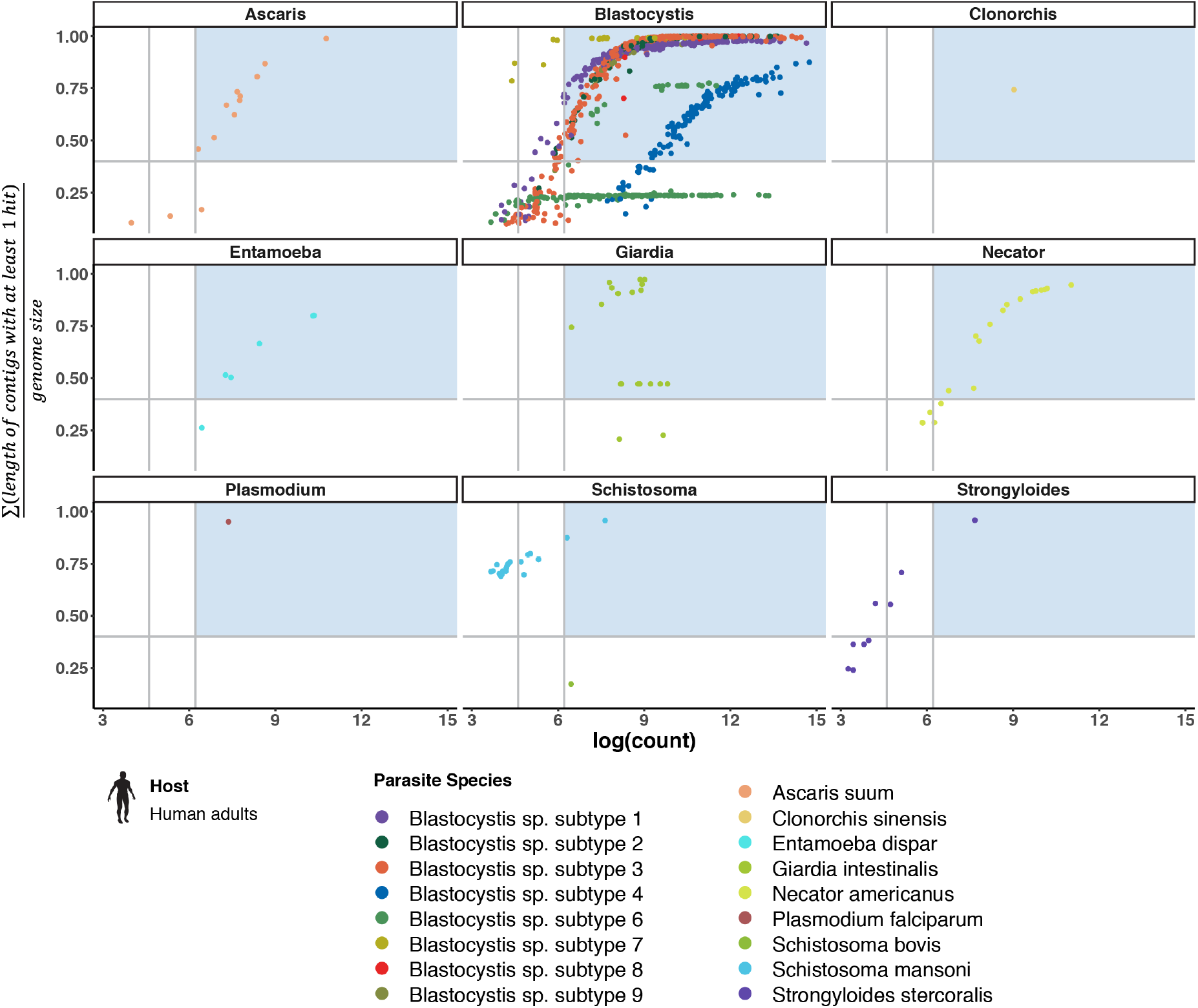
Parasites detected in human adult fecal samples. The dot plot depicts parasite-related genera in fecal samples from human adults for which at least one parasite-related species/subtype satisfied the cut-off criteria. The data points are coloured according to species. The first vertical grey line indicates the 100-read count threshold, the second line represents the selected cut-off of 500 reads observed at species level. The horizontal grey line indicates the cut-off for the sum of the length of all contigs with hits (0.4). The light blue area indicates the samples that passed the cut-off criteria.

### Blastocystis subtypes in human adult gut microbiomes

The country-level prevalence of *Blastocystis* in human adult samples ranged between 1.28 % and 100% (mean 14.3%) (Figure 5A and Table S6). Generally, the prevalence appeared higher in countries with low income as compared to high income countries (Figure 5B). The two countries with the largest samples size, USA and China, exhibited a low prevalence. Overall, the diversity of *Blastocystis* sp. subtypes was higher in high-income countries as compared to low-income countries. While all 8 detected *Blastocystis* sp. subtypes were observed in the gut microbiome of human adults from high-income countries, human adults in lower-middle and low income countries were only colonized with ST1, ST2 and ST3. In particular, *Blastocystis* sp. subtype ST4 was observed only in high-income countries like Denmark, Germany, Luxembourg, Sweden, and USA, but also in a small fraction of humans in China. A similar pattern was observed for subtype ST 6 (Figure 5B).

**Figure 5.**
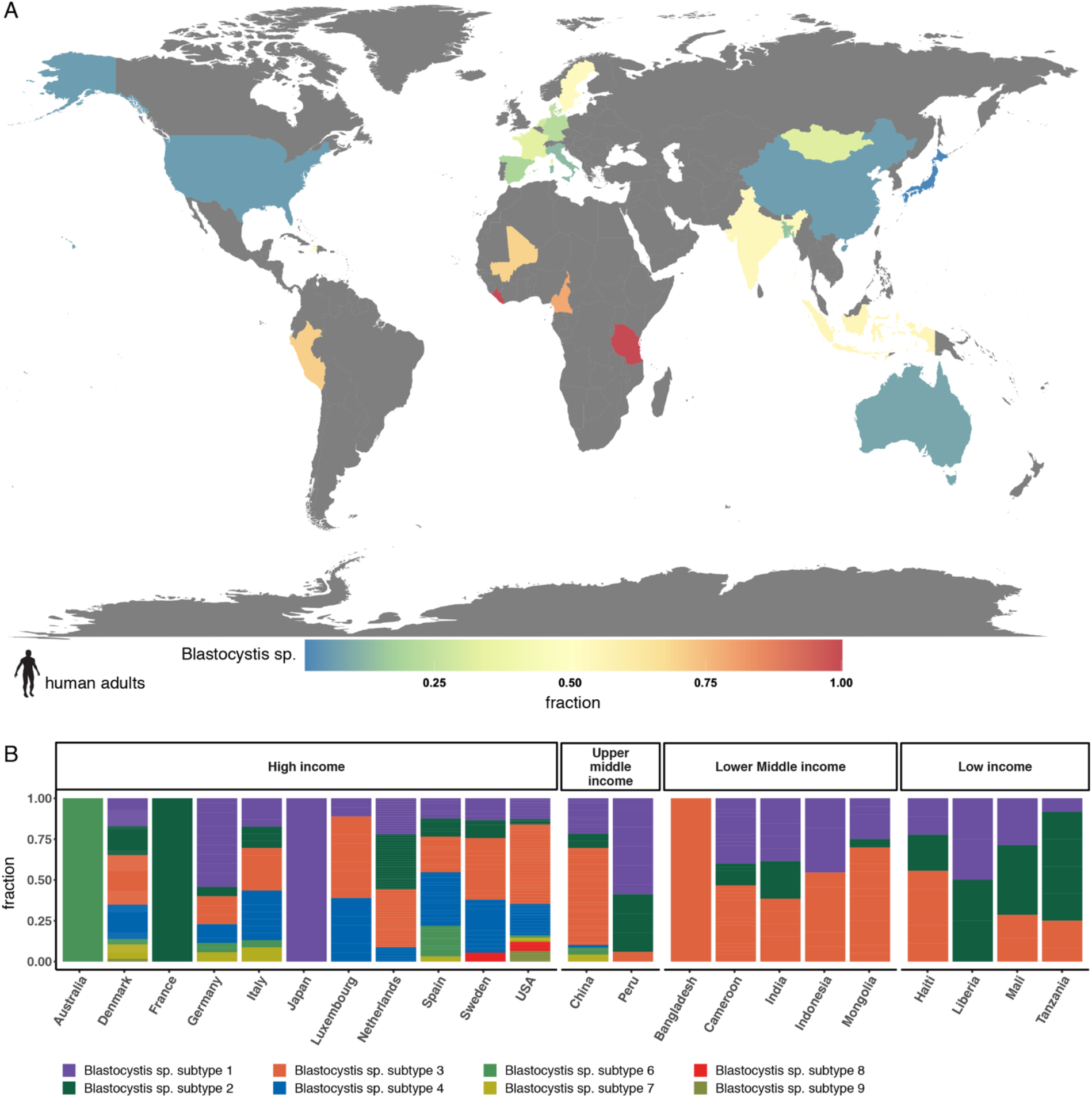
Blastocystis in human adult fecal samples. A) Geographic heatmap of country-level Blastocystis prevalence in human adult samples. The prevalence of Blastocystis was determined by metagenomic analysis and the measured values range between 1.28 % and 100%. B) The relative abundance of different *Blastocystis* sp. subtypes in different countries. The countries are grouped by income levels according to World Bank. *Blastocystis* sp. subtypes are indicated by colour.

### Parasites in human infant microbiome samples

To investigate gut parasite-colonization in early life of humans, we analysed available fecal samples from human infants from four countries (Estonia, Finland, Russia, and USA). We detected *Blastocystis* in 3.6% (n = 22) of the 609 samples from human infants (Figure S5). The range of colonization was between 0% in the USA and 12.5% in Estonia (Figure S5 and Table S7). The prevalence for *Blastocystis* in infants was 1.68% on average. Other organisms found were *Caenorhabditis elegans* and *Cryptosporidium meleagridis* in one sample respectively (Figure S5, and Tables S7 and S8).

## Discussion

Parasites belong to a variety of eukaryotic taxonomic lineages. Some parasites can be benign, co-existing within their host, and others can cause serious diseases in humans. Currently, a variety of laboratory assays exist to identify parasites in human and veterinary clinical settings. Metagenomics offers the opportunity of a one-for-all analytical approach, in which any potential pathogen, including bacteria, viruses, or parasites, could be identified based on shotgun DNA sequences generated from complex biological specimens (e.g. fecal samples) from humans and animals. While there has been great technological advancement for the identification of bacteria and viruses from complex metagenomics datasets, the identification of parasites using metagenomics is lagging behind for a variety of reasons (see Introduction). In this study, we present an analytical workflow for the identification of parasites in human and animal fecal samples. We carefully analyse metagenomic datasets from >7.000 gut microbiome samples from pigs, chicken, human adults and infants, and identify distinct parasites in these sample sets.

In pigs, *Blastocystis* was detected in 79% of all herds. The most common identified subtype was ST1. Other studies on *Blastocystis* in pigs found ST5 to be the pre-dominant subtype followed by ST1 (20–23). However, since there is no reference genome for *Blastocystis* ST5, we could not verify this using our metagenomics approach. It is possible that reads originally belonging to ST5 were assigned to similar genome sequences representing other subtypes. *Blastocystis* levels in pigs were higher in the Netherlands and Belgium compared to other European countries while Germany exhibited the lowest levels. This may be caused by different drug treatments in the respective countries (15). While there were low levels of *Blastocystis* in Germany, it was the only country where we detected *Ascaris suum*, a common pig parasite, as well as *Pristionchus entomophagus*, in one sample. However, *P. entomophagus* may have been introduced while fecal matter was collected from the pen floor as it is commonly regarded as a free-living nematode. Generally, the identification of parasites from samples that are pooled from several animals, as in this case, could pose challenges because the parasite cells are being diluted if only a fraction of animals are colonized.

In chicken, colonization with an *Eimeria* species was detected in 83% of all flocks. This is in line with other studies reporting infection rates between 80-100% based on conventional detection methods (24–26). *Eimeria acervulina* was the dominant *Eimeria* species, followed by *Eimeria maxima*, and *Eimeria tenella*. Distribution of the *Eimeria* species seems to depend on geography, as studies have reported different species to be dominant in different regions (24–26). Detection of *Eimeria* in these previous studies was performed by PCR-based methods (25, 26) and amplicon sequencing (24). *Eimeria* species cause coccidiosis, one of the diseases with the largest economic impacts on the poultry industry (27, 28). Coccidiosis is caused by different *Eimeria* species targeting different parts of the intestines and leads to reduced weight of the grown chicken even in subclinical cases (29). To our knowledge, metagenomics has never been previously used to detect *Eimeria* in chicken. We also identified *Cryptosporidium baileyi* and *Cryptosporidium meleagridis* in chicken flocks. Infection with *Cryptosporidium* causes cryptosporidiosis in a wide range of hosts, including chickens, pigs, and humans. Most common clinical signs of cryptosporidiosis are diarrhea, weight loss and dehydration (30, 31).

The differential data analysis in livestock samples revealed that *Prevotella* is positively associated with the presence of *Blastocystis* in European pig farms. This association was also found in studies of human gut microbiomes using metagenomics (32, 33). We could not find any associations of *Eimeria* with other members of the gut microbiome in chicken samples. A reason could be that the DNA from chicken samples was amplified prior to sequencing because DNA yields were not high enough originally, and in result the abundance levels might have been skewed (15).

In human adult gut samples, we observed a global distribution of *Blastocystis* ST1, ST2 and ST3 in all regions, while additional subtypes (e.g. ST4, ST6, ST7, ST8) were detected in high-income countries and China. However, the uneven distribution of samples between countries, especially between developed and underdeveloped countries, should be kept in mind. In line with our finding, Beghini and colleagues also reported that ST4 is mostly found in Europe and the USA (12). In contrast, we found that ST3 was more common in underdeveloped countries than ST2. Blastocystis subtypes ST1, ST2, and ST3 were also recently identified in human paleofeces (34), of which ST1 was most dominant. This suggests that *Blastocystis* is a common ancient colonizer of the human gut, and that *Blastocystis* colonization in general and the prevalence of distinct subtypes has changed during industrialisation. In support, it has recently been discovered that the presence of Blastocystis was associated with favourable glucose homeostasis and lower estimated visceral fat and lower BMI (35). We also identified two soil-transmitted helminths (i.e. *Ascaris suum* and *Necator americanus*) in human adult gut samples of which most were from Indonesia. These samples originated from a study about humans with helminthic infections in Indonesia (13). As these samples were biased towards specimens with helminth infection, we cannot conclude on the general prevalence of these helminths in Indonesia. Generally, we were not able to verify all results from the qPCR and Kato-Katz analysis of this study, as we could only detect *Necator sp*. in 9 of 12 samples. Furthermore, we identified 8 of 12 cases of *Ascaris* using the *Ascaris suum* reference genome, since there is no reference genome available *for Ascaris lumbricoides*. However, this might have only minimal impact on the detection, since there is evidence that *Ascaris suum* and *Ascaris lumbricoides* are a single species (36, 37). Furthermore, we did not detect *Trichuris trichiura* using our metagenomic approach in any of the samples from the study (13).

In human infants we found that *Blastocystis* presence was minimal, suggesting that colonization might happen with higher age, and/or is highly dependent on environmental factors. Detection of *Caenorhabditis elegans* in one sample may have been a false positive identification, due to undetected contaminations (38), or a potential sequencing contamination as *Caenorhabditis elegans* is a common model organism for studying human immunity (39).

Parasite identification based on metagenomics-based analyses poses multiple challenges. There are still many parasites for which no published reference genome exists, increasing the possibility of false classification or no detection at all. Challenges can also arise when working with an insufficiently large database in general. When we classified sequences using a reduced database, we encountered that reads accumulated on a set of certain contigs. Deeper investigations revealed that those contigs contained ribosomal DNA. Ribosomal DNA is a potential source for misclassification especially when the actual source organism (plant genomes, in this particular case) is not present in the database and the taxonomic tree is only sparsely covered since ribosomal DNA is highly conserved even between distant species.

Human fecal samples are one of the most studied sample types in metagenomic sequencing. However, no standardized DNA extraction method is being used across studies. It has previously shown that DNA isolation procedures can have an impact on inferred microbiome composition (16). For the human gut microbiome samples from different studies that we analysed here, a range of DNA extraction methods were being applied, often with modifications such as an additional lysis step (Figure S6, and Table S9). Many metagenomic DNA extraction protocols are optimized for bacterial extraction and were not assessed regarding parasite identification. This could lead to that certain parasite stages, such as oocysts or helminth eggs, are not being disrupted and their DNA not made accessible for sequencing. This makes prevalence assessments for parasites obtained from current metagenomic studies less reliable. For instance, we see large variations in *Blastocystis* prevalence for the different extraction methods within the same country: when examining samples from Germany, the fraction of *Blastocystis*-positive samples is similar for 2 of the 3 methods, however no *Blastocystis* are detected when using the PSP Spin Stool DNA plus kit from Invitek. Furthermore, when using the QIAamp DNA Stool Mini kit (Qiagen), an additional beat-beating step seems necessary to increase the chance for detection of *Blastocystis* (Figure S6, and Table S9). We also discovered in a recent study on microbiome sample storage conditions that parasite oocysts from *Cryptosporidium parvum* were better detected upon several freeze-thaw cycles of the sample (40).

While challenges remain, we believe that metagenomics has the potential to serve as a robust method for parasite detection for a wide range of samples. Current detection methods often rely on microscopic examination of the sample or specific PCR. Metagenomics-based analyses could facilitate a faster and more convenient way of detecting parasites in humans and animals. The metagenomic approach could serve as a one-for-all untargeted approach for pathogen identification, including bacteria, viruses, and parasites. The results from our study appear promising, and additional research is needed to verify the reliability of the metagenomics-based approach. For follow-up studies we suggest performing conventional and metagenomics-based methods on the same sample set, including suitable positive and negative controls.

## Material & Methods

### Sample acquisition and pre-processing

Livestock metagenomic shotgun sequencing reads from fecal samples from pig and chicken farms were originally collected for the EFFORT study on antimicrobial resistance in nine European countries by Munk and colleagues (15). The fecal samples were extracted using a modified QIAamp Fast DNA Stool Mini Kit protocol (16), and sequenced on a HiSeq3000 (Illumina) platform. The resulting sequence files were downloaded from NCBI and pre-processed as described in the original publication. This resulted in 181 samples from pig herds and 178 metagenomic samples from chicken flocks. Each sample represented a pooled sample originating from 25 fresh undisturbed pen floor fecal samples, respectively (17).

Human adult metagenomic datasets from stool samples with more than 10 million paired-end reads were downloaded from ENA, and they originated from 51 microbiome projects (Table S10). This resulted in 6.152 samples from 22 different countries. Additionally, we downloaded 609 publicly available metagenomic datasets originating from human infant fecal samples from 4 different countries and 6 different microbiome projects (Table S10).

### Database

The database for metagenomic classification with kraken2 v2.0.8-beta (18) (see below) consisted of 162.318 genomes of which 530 were parasites (protozoa, flatworms, roundworms) or genomes close in the taxonomic tree of known parasites (Table S11). Parasite and parasite-related genomes, obtained from species close to a parasite in the taxonomic tree, were selected from a pool of 926 protozoan, flatworm, and roundworm genomes downloaded from GenBank. Since assemblies of parasites are often of lower quality compared to bacterial genomes, a subset of samples acquired from the EFFORT study was classified using kraken2 with a database containing the 926 parasites and closely related genomes. All contigs with hits were selected to be analyzed by querying them against the NCBI ‘nt’ databases as described in our previous study (6), with the modification that a contig would be flagged as a potential contamination if there was a match against an organism outside of its own genus with a coverage of above 30%. Genomes were then manually selected, up to three for each species, based on assembly statistics and the number of potentially contaminated contigs. Further, the potentially contaminated contigs were removed from the assemblies. Since we previously showed that contaminated contigs are mostly below 5.000 bases long (6), we removed all sequences shorter than 1.000 bases. This threshold was selected because removing larger contigs may in some cases mean that large parts of a whole assembly are removed. To increase the resolution for the parasite genomes we introduced two new taxonomic levels: assembly and contig. If there are multiple assemblies available for one species, we created a node in the taxonomic tree for each assembly (below the species level). Additionally, each contig was attached to the assembly node as a leaf. If there was only one assembly available, we connected the contigs directly to the species node.

### Metagenomic classification and statistical data analysis

Metagenomic classification was performed using kraken2. Kraken2 is a k-mer based method that uses the lowest-common-ancestor (LCA) approach to assign reads to a taxonomic identifier. If a read can be assigned to two different organisms equally well, the read will be assigned to the lowest-common-ancestor of both organisms in the NCBI taxonomy.

For statistical analysis, kraken2 results with confidence parameter set to 0.15 were used for livestock and human samples. To reduce the dimensions of the count matrix, all counts below the species level were moved up to species level. Species level counts represented with fewer than 100 reads were moved up in the taxonomic tree to further increase confidence in our results.

Parasites and parasite-related organisms were only reported as a true hit if the following quality cut-offs were met:

1. At least 500 hits were observed at species level
2. 5% of all contigs in the assembly had at least one hit
3. 40% of all bases in the assembly are represented by contigs with at least one hit

Since we observed many hits to *Blastocystis* in pig herds and *Eimeria* in chicken flocks we created two groups in each sample type: a negative group (non-colonized) and a positive (colonized) group. Samples were defined as positive (colonized) based on the above-described quality cut-offs.

Center log-ratio (clr) transformation of the count matrix and compositional data analysis was performed using the aldex2 package version 1.20 (19) for R version 4.02. The number of Monte Carlo samples used to estimate the underlying distributions was set to 128 and features to use as the denominator for calculating the geometric mean was set to “all”. Statistically significant differences in organism abundances between countries and between groups colonized with *Blastocystis* or *Eimeria*, respectively, were determined using the effect size (greater than 1 or smaller than -1) as well as Benjamini-Hochberg corrected p-values smaller than 0.005 for the Welch’s t-test and Wilcoxon Rank Sum test from the aldex2 output.

## Acknowledgements

This work was in parts supported by the European Union’s Framework Programme for Research and Innovation, Horizon2020 (643476). The funders had no role in study design, data collection and interpretation, or the decision to submit the work for publication.

