## Supplementary Figures for "Detection of Parasites in Microbiomes using Metagenomics"

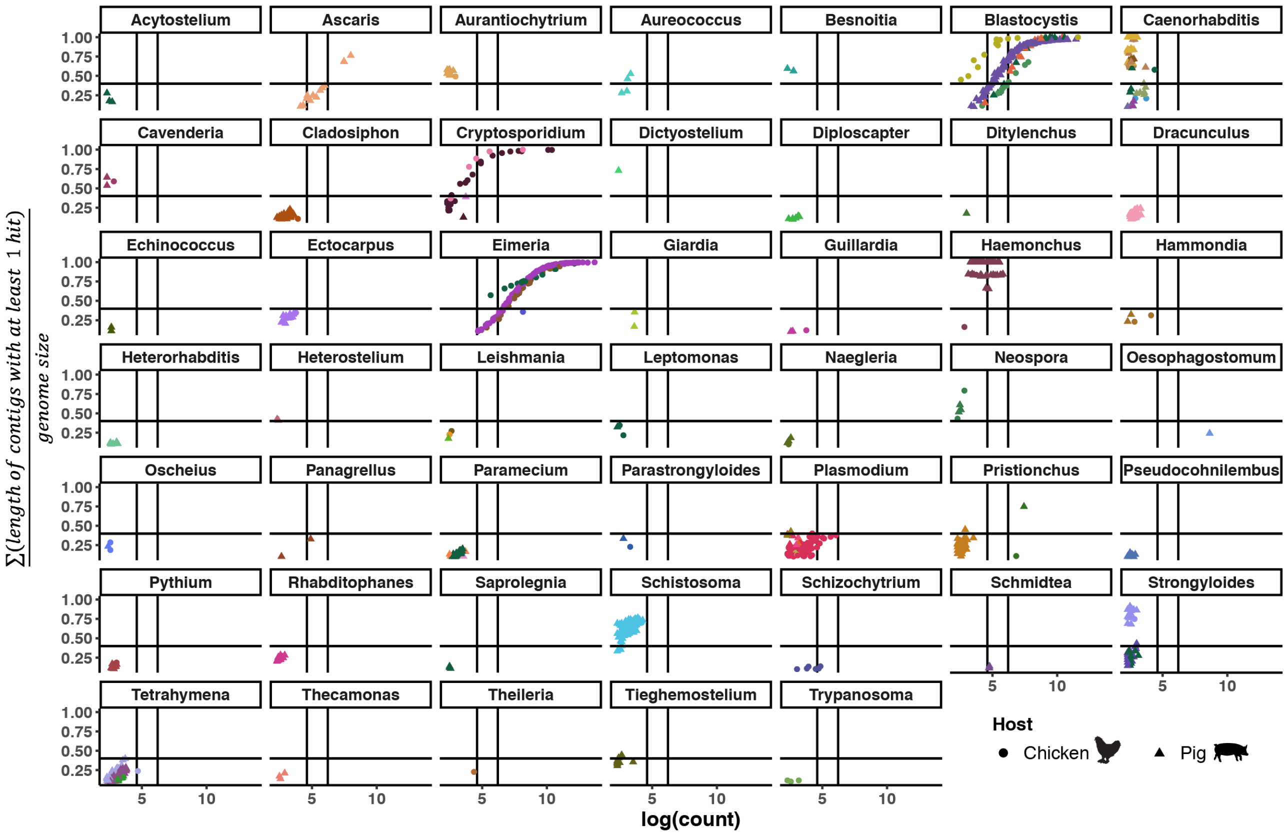


**Figure S1. All parasites detected in livestock samples.** The dot plot depicts parasite-related genera in livestock samples for which at least in one sample more than 1 counts per million (cpm) where detected. The data points are coloured according to species and indicated by the host the host they originate from (dot: chicken; triangle: pig). The first vertical line indicates the 100-read count threshold, the second line represents the selected cut-off of 500 reads observed at species level. The horizontal line indicates the cut-off for the sum of the length of all contigs with hits (0.4).

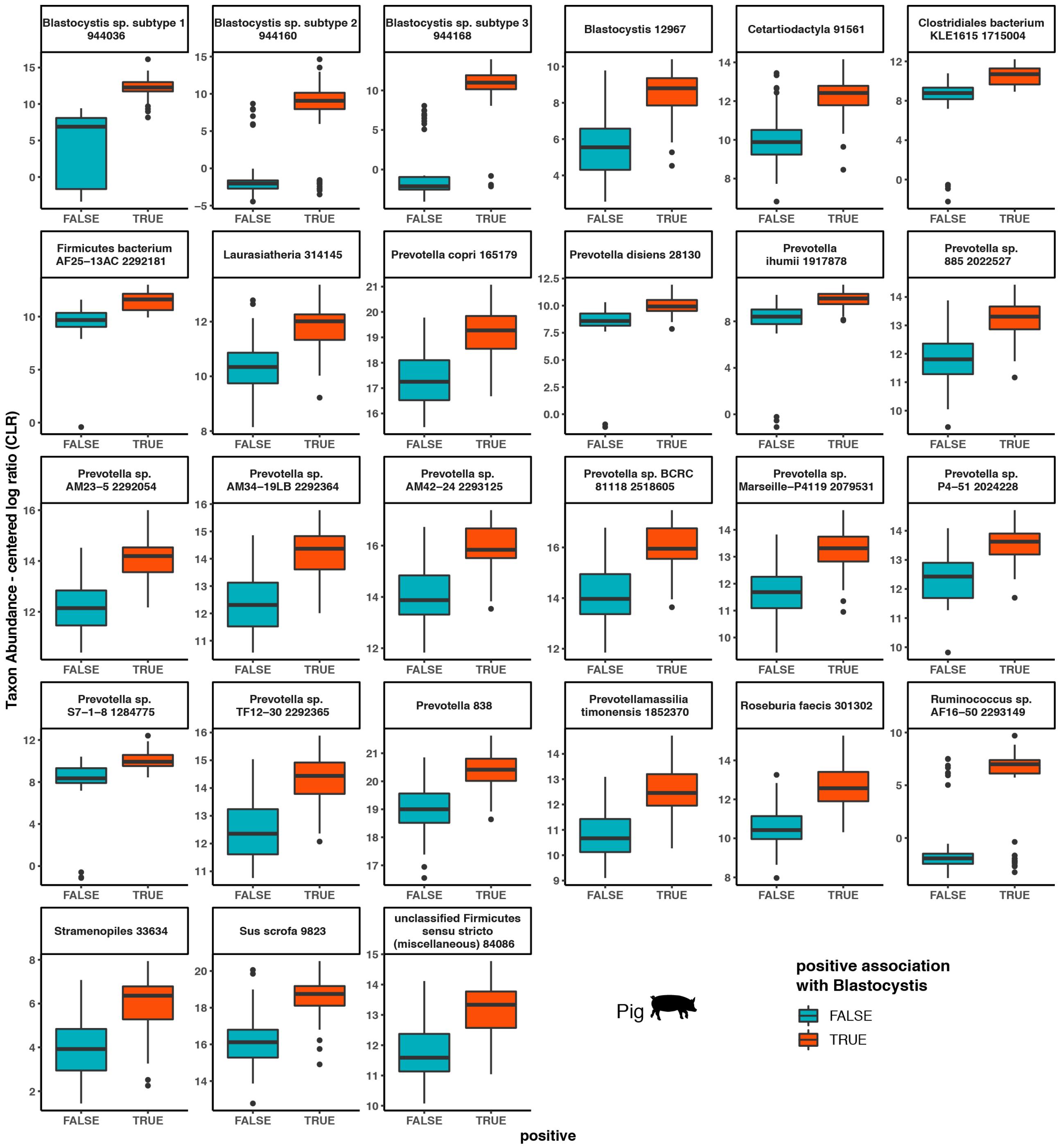


**Figure S2. Association between Blastocystis and other members of the gut microbiome.** Differential data analysis of pig gut microbiome taxa in relation to samples with low/no Blastocystis presence (teal) against samples with high Blastocystis presence (orange). The effect size had to be > 1 or < -1, and a Benjamini-Hochberg corrected p-value smaller than 0.005 for the Welch's t-test and Wilcoxon Rank Sum test from the aldex2 output.

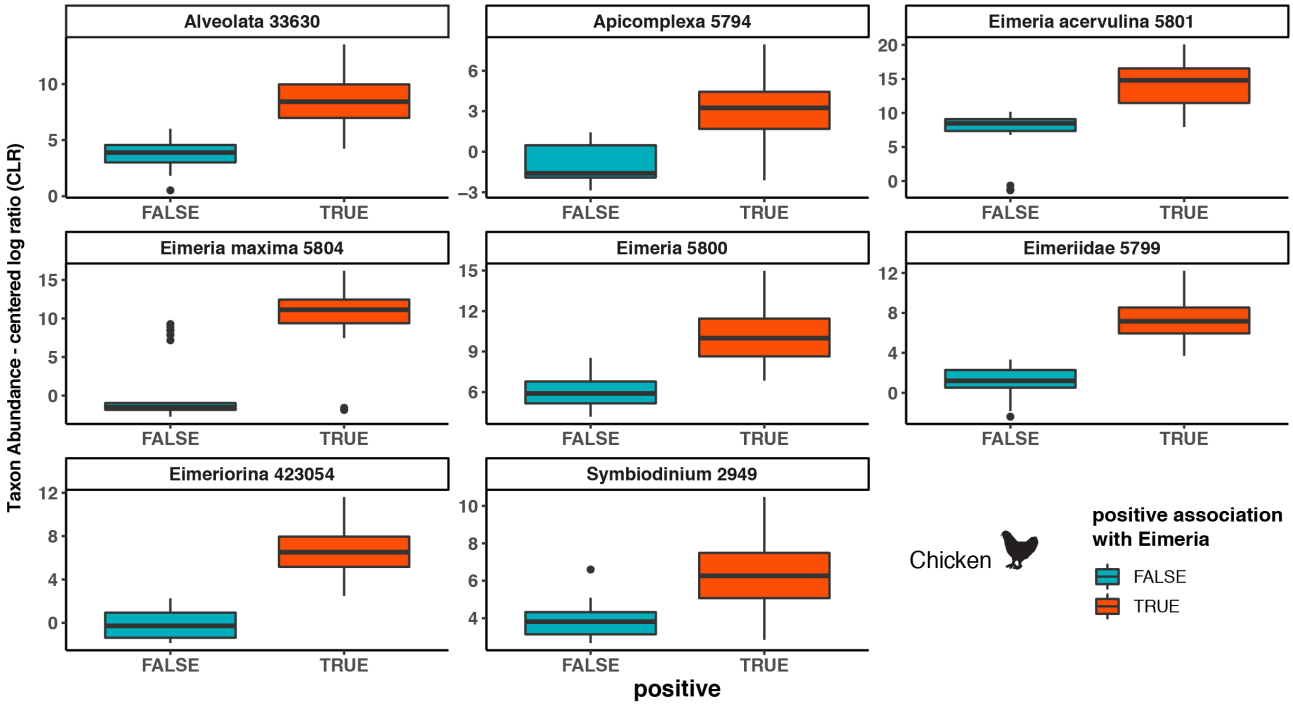


**Figure S3. Association between Eimeria and other members of the gut microbiome.** Differential data analysis of chicken gut microbiome taxa in relation to samples with low/no Eimeria presence (teal) against samples with high Eimeria presence (orange). The effect size had to be > 1 or < -1, and a Benjamini-Hochberg corrected p-value smaller than 0.005 for the Welch's t-test and Wilcoxon Rank Sum test from the aldex2 output.

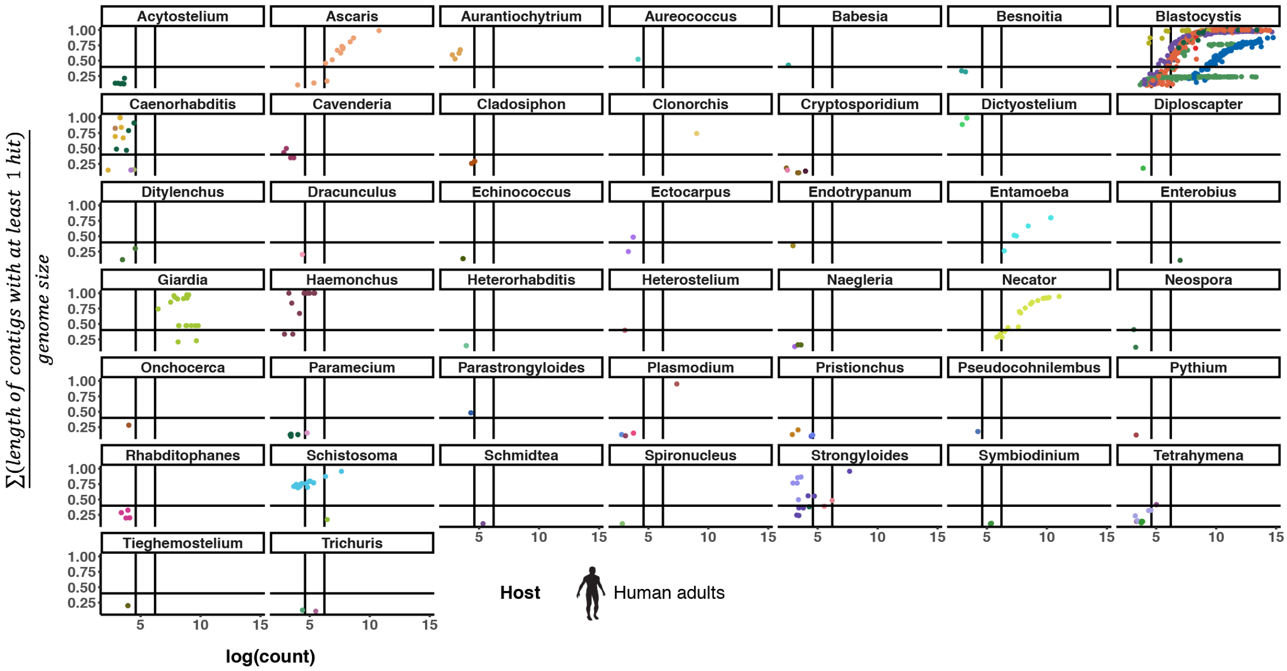


**Figure S4. All parasites detected in human adult gut samples.** Dot plot of parasites in adult human samples where at least in one sample more than 1 counts per million where detected. Data points are colored according to species. The first vertical line indicates the 100-count threshold, and the second the selected cut-off of 500. The horizontal line indicates the cut-off for the sum of the length of all contigs with hits (0.4). Nine taxa passed the cut-off criteria (section 2.3), and these are displayed in Figure 4.

**Detection of Parasites in Microbiomes using Metagenomics**
Philipp Kirstahler, Frank M. Aarestrup, and Sünje Johanna Pamp

A

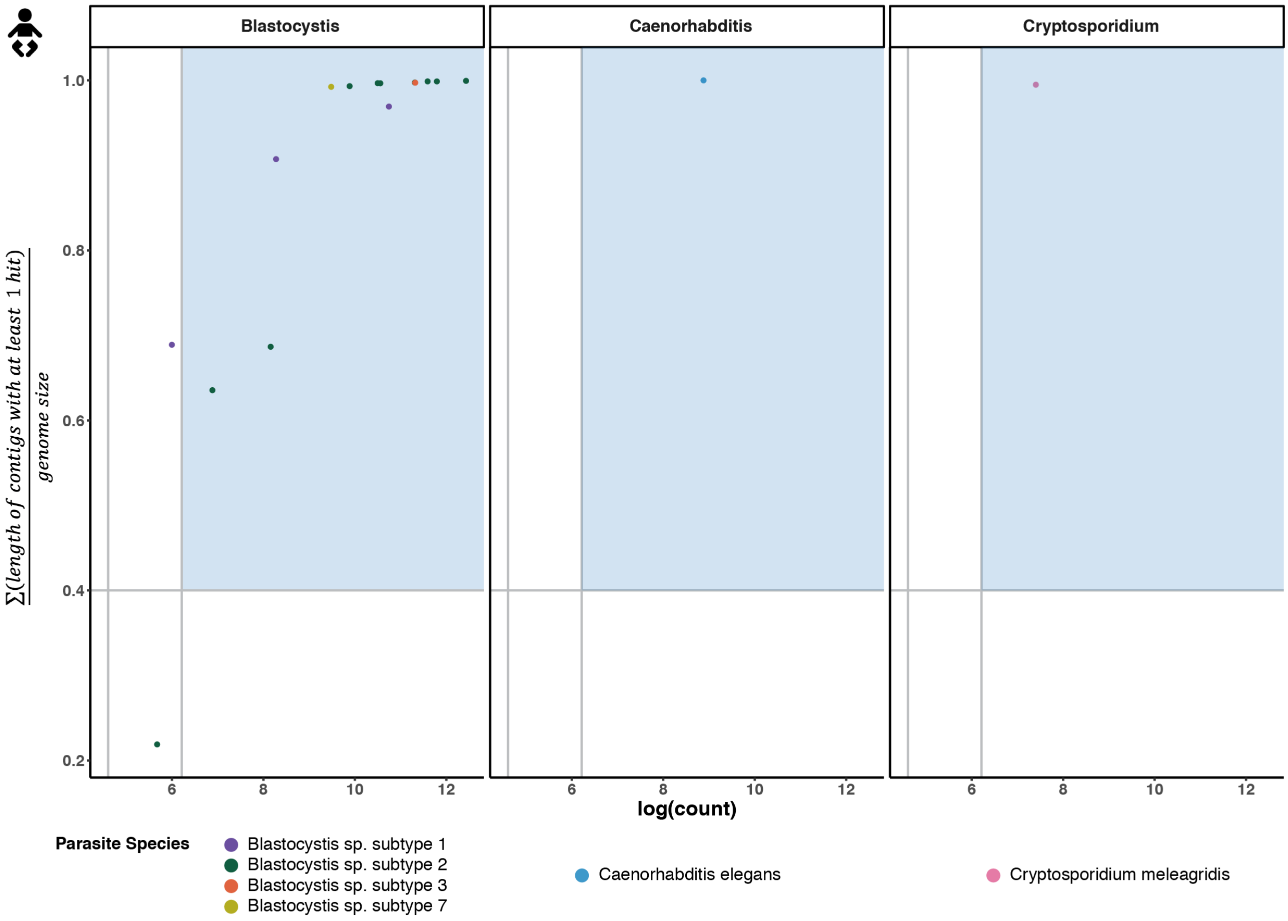


B

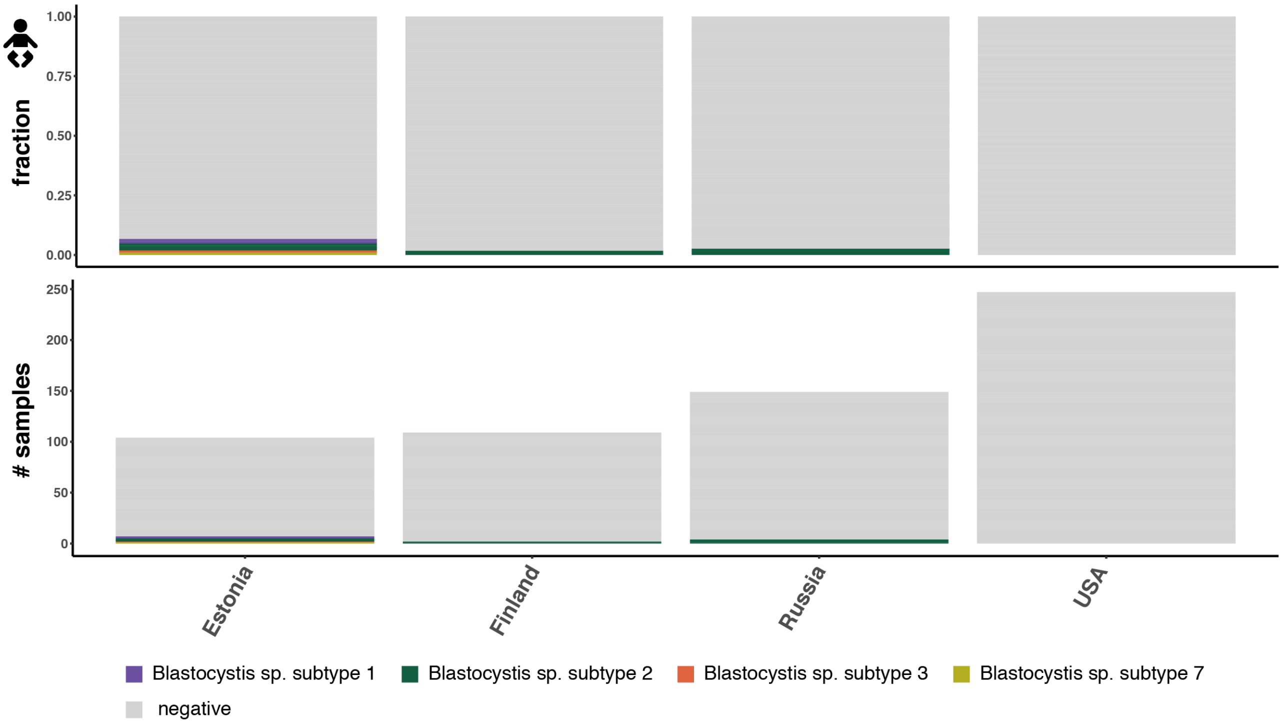

**Figure S5. Parasites detected in human infant gut samples.** A) Dot plot of parasites in human infant fecal samples where at least one sample satisfied the cut-off criteria described in the method section. The data points are colored according to parasite species. The first vertical line indicates the 100-count threshold, the second line the selected cut-off of 500. The horizontal line indicates the cut-off for the sum of the length of all contigs with hits (0.4). B) Parasites detected in human infant samples based on the country they originate from. The bar plot at the top indicates the faction of samples, and the bar plot at the bottom indicated to total number of samples per country.

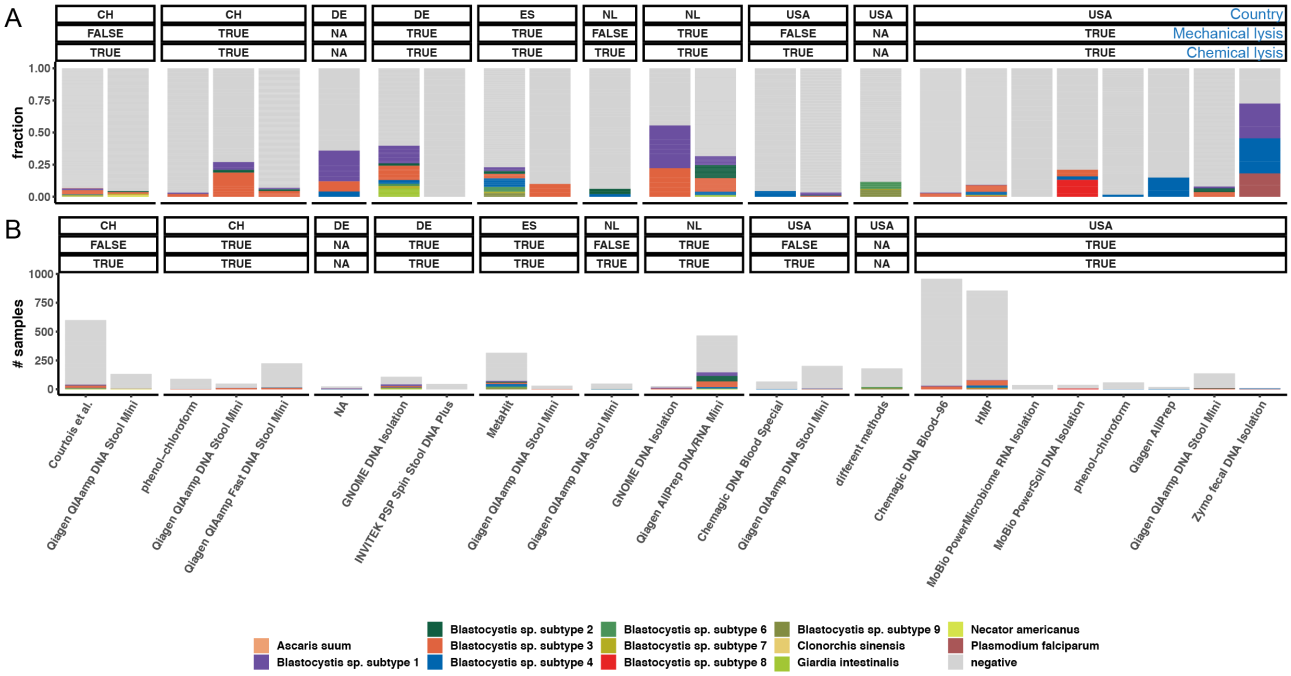


**Figure S6. DNA isolation procedures**. These bar plot displays the DNA isolation conditions for the human gut microbiome studies whose samples were analyzed as part of this study. The samples are grouped by DNA extraction method, country, and cell lysis steps (mechanical and chemical). A) Displays the fraction of samples for which parasites were detected. B) Displays the absolute number of samples containing parasites. For some aspects, no information could be extracted from the literature, and these are marked with NA.
